# Characteristics of spatial summation in the magnocellular, parvocellular, and koniocellular pathways

**DOI:** 10.1101/2024.10.11.617932

**Authors:** Christopher S. Wu, Daniel R. Coates

## Abstract

In this study, we characterize the spatial summation properties of targeted magnocellular, parvocellular, and koniocellular pathways within the central 20° of visual field using chromatic transformations in DKL color space. For the magnocellular and koniocellular conditions, critical areas of complete spatial summation were found for all eccentricities. For the parvocellular conditions, complete spatial summation was absent within the stimulus size ranges tested. We also describe an anatomically and physiologically motivated model of receptive field pooling using probability summation. Model simulations suggest that the critical area of summation can be explained by the dendritic field size of underlying retinal ganglion cells, corroborating our psychophysical data.

## 1. INTRODUCTION

The measurement of localized detection sensitivity is widely accepted as the clinical standard for assessing visual function in the context of certain ocular and neurological conditions. Simple non-invasive tests such as perimetry measure contrast sensitivity loss at discrete visual field locations that may occur as a result of disease pathology. Despite its ubiquity however, the relationship between structural changes in the visual system and functional sensitivity is not fully understood. In this study, we characterize the effects of stimulus size on the detection of achromatic and chromatic stimuli across the visual field. We also provide an anatomically and physiologically motivated model to describe changes in visual sensitivity in terms of receptive field morphology and neural pooling characteristics. This is explored in the context of the distinct magnocellular, parvocellular, and koniocellular pathways originating in the retina. Understanding how detection thresholds change in this context may reveal underlying neural mechanisms and provide methods to improve test diagnosticity.

Visual spatial summation describes contrast detection thresholds as a function of stimulus size. In the spatial summation of achromatic stimuli, increases in stimulus size result in proportional improvements to detection thresholds following Ricco’s Law [1]. This complete summation area presents with a slope of −1 when plotted as log thresholds as a function of log stimulus area. Beyond a certain threshold size however, known as Ricco’s Area or the critical area (A_C_), further increases in stimulus size result in a reduced improvement in sensitivity. This incomplete or partial summation area presents with a shallower slope between 0 and −1. There has been much interest in the anatomical or physiological correlates of the A_C_.

The reduction in summation ability beyond a critical size has naturally been attributed to the spatial limit of some neural mechanism in the visual system. Early electrophysiological studies discovered areas of complete spatial summation within the receptive fields of retinal ganglion cell (RGC) units [2, 3, 4]. It was hypothesized that these same receptive fields were responsible for the psychophysical A_C_, although inconsistencies in how the size of these receptive fields change with eccentricity led to the proposal of further retinal or cortical contributions [5, 6]. In response to the relatively poor correspondence between changes in RGC size and the psychophysical A_C_ with retinal eccentricity, others have considered instead the overlap of receptive fields [7, 8], or their density [9, 10, 11, 12, 13].

While evidence towards a retinal locus for spatial summation is compelling, recent studies on disease models suggest potential cortical contributions. For example, the enlargement of the A_C_ in glaucoma [11] has been proposed to contradict a retinal basis due to the shrinkage of RGC axons and dendritic trees [14]. Likewise, the A_C_ was found to be larger in amblyopes [15] despite their apparent normal retinal function and architecture [16, 17, 18]. Neural pooling models presuming a cortical component were capable of reproducing the spatial summation function as well [19, 20, 21]. Naturally, for visual information to influence decision-making such as in psychophysical tasks, there must be some cortical component, however, the question lies in the anatomical or physiological basis of the A_C_ and other features of the spatial summation function.

To explore this question, we characterize the spatial summation properties of chromatically targeted magnocellular, parvocellular, and koniocellular visual processing pathways as a function of retinal eccentricity. This is done within the axes of Derrington, Krauskopf, and Lennie (DKL) color space, designed to selectively target theorized parallel processing pathways [22]. While the effects of eccentricity on the achromatic spatial summation function have been well documented [6, 23], their potential association with RGCs is hindered by the ambiguity in contributing cell type(s). Efforts have also been made to measure spatial summation functions within cone specific mechanisms [9, 10, 24], although their relationship with specific RGC types is complex. We are only aware of one study that has characterized the spatial summation of post-receptoral visual processing pathways as defined by the cardinal axes of DKL color space [25], however, in this study, comparable achromatic data is absent and only a single eccentricity was tested.

By bridging the gap between functional sensitivity and some anatomical or physiological characteristic of the visual system, we hope to elucidate basic mechanisms of visual processing, and contribute to the development of more sensitive and disease specific clinical tests [11, 23, 26]. In describing a simplified model of receptive field pooling to account for the spatial summation function and how it changes as a function of visual pathway and retinal eccentricity, we offer a potential explanation for the psychophysical critical area and provide the framework for the development of improved perimetric stimuli.

## 2. METHODS

### 2.1 Subjects

Four subjects (mean age: 26.75 years, range: 24 – 30) were recruited for participation in this study from the University of Houston. All subjects were examined for normal ocular health and history to exclude factors that may affect contrast sensitivity. Normal color vision was verified and assessed with the Hardy Rand Rittler (HRR) pseudoisochromatic plates and the Farnsworth D-15 color test. Participants were either emmetropic or optically corrected, and special care was taken to ensure spectacle corrections were free from color blocking filters and tints. One of the participants was an author of the study, while the others were naïve to the purposes of the experiment. The experimental protocol was approved by the Institutional Review Board of the University of Houston, and informed consent was obtained for all participants. The experimental procedures were conducted in adherence to the Declaration of Helsinki.

### 2.2 Apparatus

The psychophysical experiment was back-projected on a FilmScreen 150 projector screen (Stewart Filmscreen) using a DLP LED PROPixx projector system (VPixx Technologies Inc) at a resolution of 1920 x 1080 pixels, a refresh rate of 120 Hz, and a color depth of 8-bits. The experiment was limited to a 1000 x 1000 pixels square centered on the projector screen. Subjects sat 50 cm from the display, allowing the display area to subtend 50° in the test eye, and were stabilized with a forehead and chin rest. Overall height was adjusted to ensure subjects were eye level with the central fixation cross.

The overall luminance of the display was measured with a Konica Minolta LS160 luminance meter (Konica Minolta), to ensure a linear gamma profile. The chromaticity and color calibrations were measured and confirmed with a BLACK-Comet spectrometer (StellarNet Inc).

### 2.3 Stimuli

The experimental protocol was developed in JavaScript with the jsPsych library, with a custom graphical shader overlay using the OpenGL Shading Language (GLSL). Color space conversion calculations were performed in GLSL to allow parallelized processing and pre-render manipulation of pixels. Temporal dithering of pixels was implemented through the noisy-bit method to support luminance resolution finer than 8-bits [27].

Target stimuli were defined as increments along the cardinal axes of DKL color space from a background centered at the white point (International Commission on Illumination (CIE) 1931 coordinates: 0.327, 0.335), with an overall mean luminance of 10 cd/m^2^. A total of 5 color conditions were tested: +(L+M) (achromatic magnocellular, targeting the parasol RGCs), +(L-M) and -(L-M) (chromatic red/ green parvocellular, targeting the midget RGCs), and +S-(L+M) and -S-(L+M) (chromatic blue/ yellow koniocellular, targeting the small bistratified RGCs) (Figure 1). Color space conversions were accomplished following Brainard [28]. The stimuli were solid circular patches superimposed on the background. In chromatic conditions, luminance noise was added to potential target locations to minimize non-chromatic cues for detection.

**Fig. 1.**
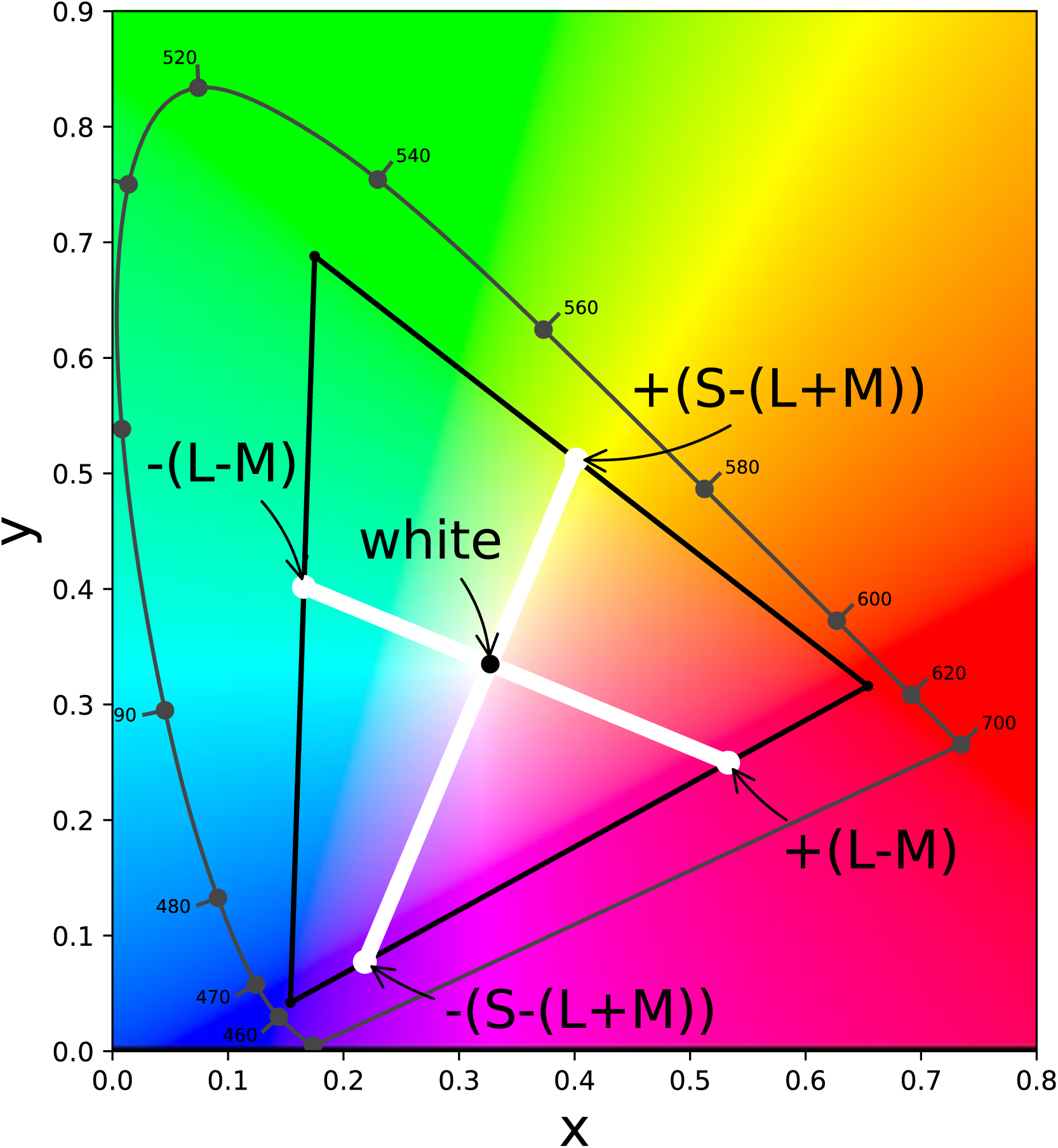
CIE 1938 diagram with the measured gamut of the display (triangle). White lines indicate the chromaticities of the cardinal axes of DKL color space, with their intersection indicating the white point of the background. The 2D space shown would represent an isoluminant plane, with the achromatic condition defined by a third axes coming out of the page at the white point.

**Fig. 2.**
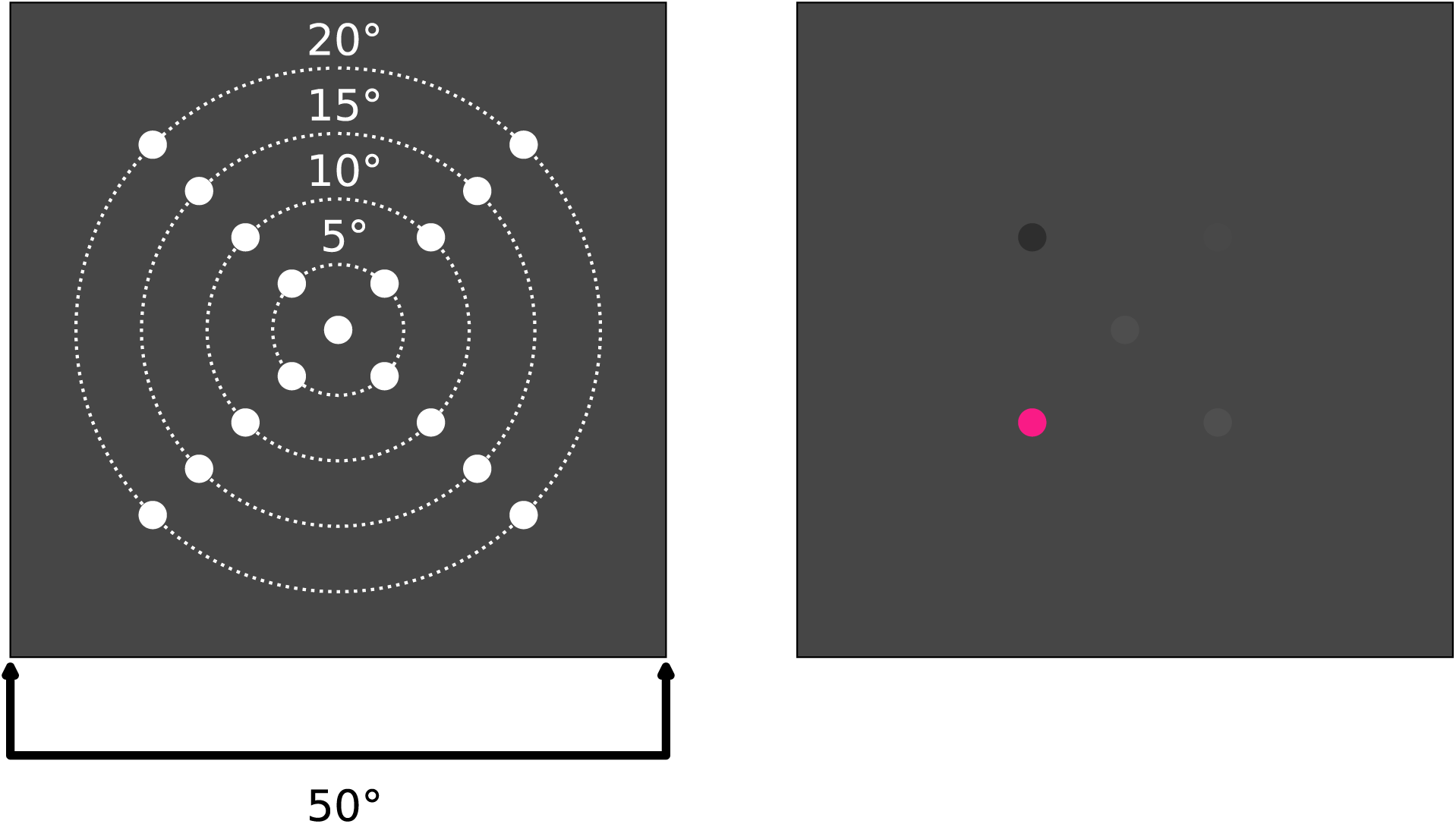
**(left).** Example experimental display with all stimulus positions shown at a set stimulus size and maximum stimulus contrast. **(right).** In each experimental block, the foveal and four radial positions at a set eccentricity are tested. Shown is red condition, 10° eccentricity. Note random luminance noise at non-target locations.

**Fig. 3.**
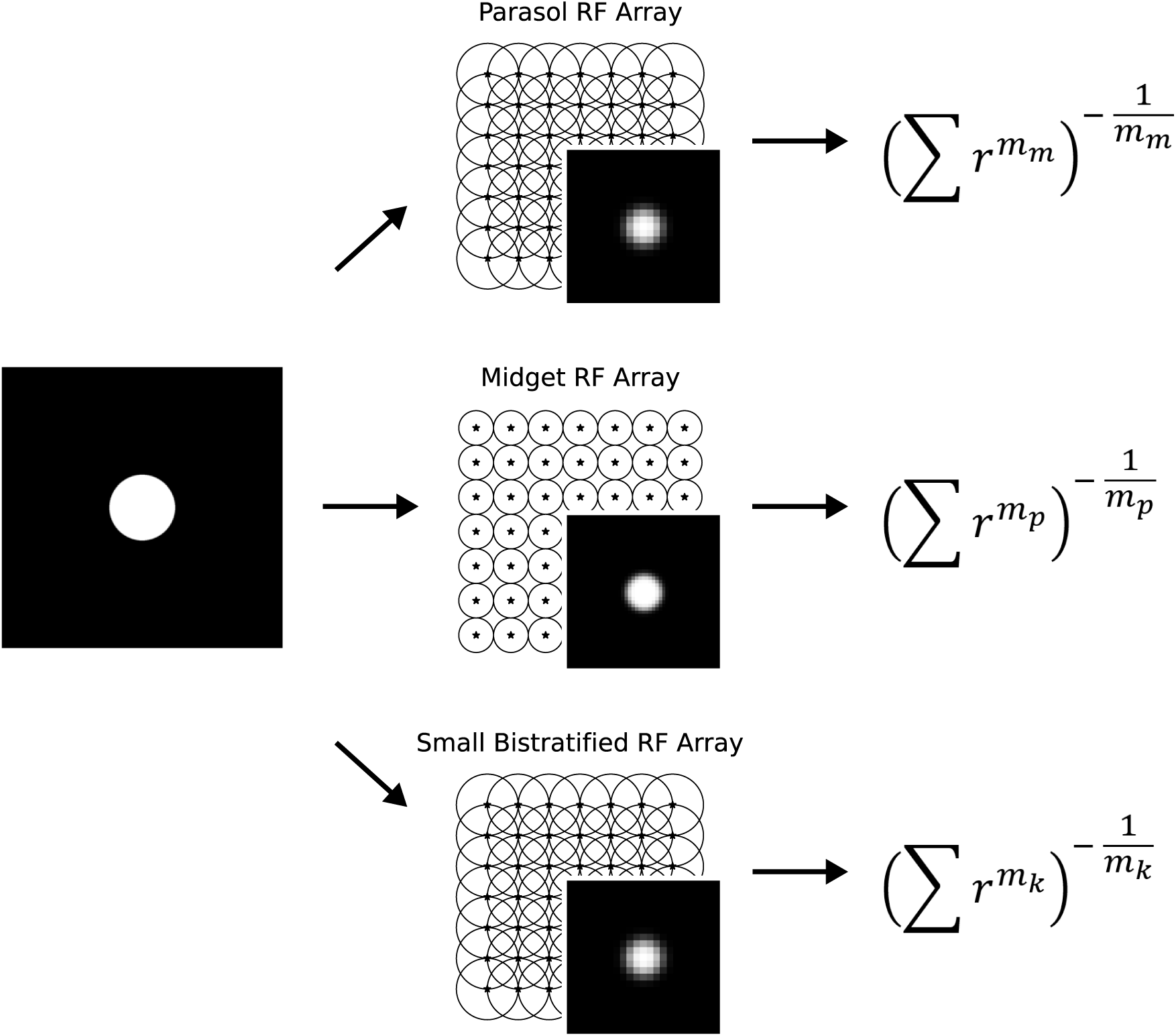
Model schematic showing progression from input image through modelled parallel processing pathways to the calculation of relative detection thresholds through probability summation. On the left, the initial stimulus is shown. In the middle column, the cone image is downsampled by the underlying retinal ganglion cell receptive field array based on anatomically-inspired sizes and overlap factors. On the right, each RGC output is pooled through probability summation, with pathway specific *m* exponent, yielding contrast detection thresholds. To simulate the whole spatial summation function, stimuli of multiple sizes are passed through the model (see Results).

### 2.4 Procedure / Paradigm

Each experimental block consisted of a single color direction, at a single eccentricity, and a specific stimulus size. In a given block, stimuli were presented at the primary oblique meridians of the visual field, along the 45°, 135°, 225°, and 315° meridians, representing the visual quadrants. Oblique meridians were chosen for this task to minimize differences in visual sensitivity with polar angle [29]. A foveal test spot matching the size of the eccentric targets was also presented to maintain fixation and attention during each block. For each color condition, up to eight stimulus sizes were tested at 5° increments up to 20° retinal eccentricity.

Contrast detection thresholds were obtained for the five test positions using multiple interleaved adaptive one-up one-down staircase procedures for each of the positions. The initial staircase step size of 1.26 log units, were reduced following each successive reversal to a final step size of 1.13 log units. The completion of each staircase required at least six reversals, while thresholds were calculated from the average of all but the first two reversals. The staircases were conducted as a function of linear distance from the white point in DKL color space along the cardinal axes of DKL color space (Figure 1). Chromatic contrasts were defined along either the incremental or decrement of the axes, decreasing in saturation (contrast) towards the white point. Reported thresholds below reflect conversion of these measurements to LMS cone contrast [28, 30]. This is a standardized characterization of input contrasts in terms of cone excitation between all conditions and is not meant to isolate the effects of cones.

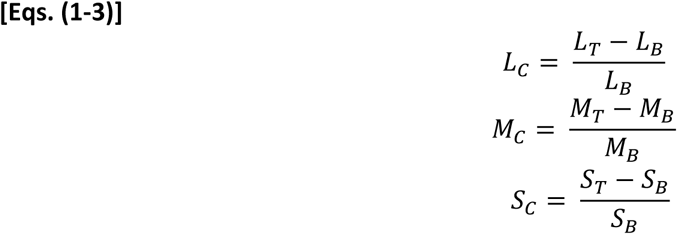

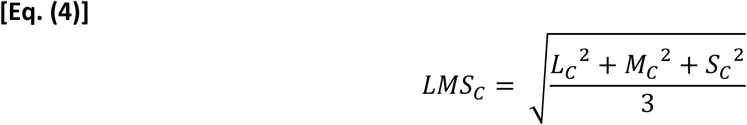

Here, individual contrasts within the L, M, and S dimensions are calculated as a Weber fraction against the background, then combined by taking the square root of the sum of squares divided by 3.

During each trial, the four eccentric stimulus locations along with the foveal target randomly change in luminance at randomly selected intervals. The intensity of this luminance noise was randomly selected within a ±10% range of the display’s background (i.e. between 9-11 cd/m^2^), while the duration was randomly selected between 250-350 ms. The target onset was randomized between 1000-1400 ms after the start of each trial, and the target for detection was presented for a duration of 300 ms. Subjects were given 3000 ms to respond after stimulus offset. A failure to respond was treated as incorrect. Unique feedback tones were given to indicate a correct or incorrect response, and to indicate start of each trial.

Prior to each experimentation period, subjects were adapted to the mean luminance of the display for five minutes. No more than 8 blocks were tested for a subject in a given period, and care was taken to prevent multiple color paradigms from being tested on the same day. During each trial, subjects were tasked with detecting and identifying the location of the presented stimuli. Responses were obtained using the corresponding oblique keys on the number pad of a keyboard (1, 3, 7, 9), with the central key (5) used for the foveal test spot. Breaks were allowed between test block, with an additional adaptation period prior to resuming.

### 2.5 Analysis/ Mathematical Model

To fit our data, we used a single continuous hinged function defined below [31].

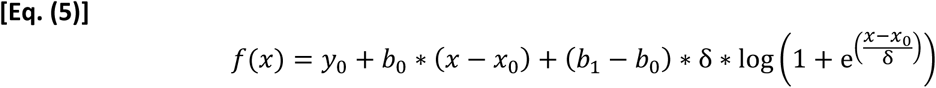

Here, five parameters are used to define the coordinates of a hinge point (*x*_0_, *y*_0_), the slopes of the two line segments that extend from this hinge point (*b*_0_, *b*_1_), and the sharpness of the hinge (δ). Following convention, the slope of the initial segment (*b*_0_) representing complete spatial summation was constrained to −1. In addition, to better characterize the hinge, the δ parameter was constrained to 0.01. Further constraints were implemented on *b*_1_ within each chromatic condition based on initial estimates of the slope of partial summation. Special care was taken in constraint selection to avoid biases from a floor or ceiling effect, and to improve bootstrap fitting described below.

Data sets were analyzed separately for each individual subject, color modulation direction, and retinal eccentricity. Each set containing thresholds for all sizes comprising the spatial summation function for the four retinal meridians was bootstrapped, randomly resampling among the four test positions. The continuous hinged function was fit to each bootstrap using the non-linear least squares method. Reported function parameters are the average of 1000 bootstrap iterations.

### 2.6 Retinal/Cortical RF Model

To better understand how the neural and morphological properties of underlying mechanisms within the targeted pathways may contribute to the psychophysical spatial summation function, we describe a basic model of neural pooling. The goal in developing this model is to provide a simple explanation for features of the spatial summation function, and ultimately bridge functional measures of sensitivity to anatomical and physiological changes in disease.

At its core, this model utilizes a cascade of convolutional filtering layers and downsampling or pooling layers to simulate the progression of information through the visual system [32]. The final output of the model in terms of relative detection thresholds is calculated through probability summation [19, 20, 21, 33, 34, 35]. To understand the effects of receptive field size and the probability summation exponent, stimuli of multiple sizes are simulated through various combinations of parameters based on reported measures. The resultant simulated functions are then fit by the continuous hinged function described above and compared to the human psychophysical functions.

At the initial stage, the input image or stimulus is sampled based on the size and resolution limit of the underlying cones photoreceptors. At this level, the pixels comprising the image are scaled according to cone densities as a function of retinal eccentricity [36, 37]. Here, cone responses to the maximum stimulus contrast are calculated to provide relative output sensitivies. For simplicity, the selectivity of cone types were not considered, and each simulated cone contributes equally to underlying retinal ganglion cells regardless of class.

The cone image is subsequently passed through the convolutional filtering layer of the model which represents pooling by underlying RGCs. In this layer, a Gaussian filter that characterizes the sensitivity profile of a single retinal ganglion cell receptive field center [4, 5, 38, 39, 40] is convolved with the cone image. Thus, the cone image is downsampled to the resolution of the RGC array. For simplicity, RGC sensitivity is invariant to retinal eccentricity, rather only the size of their receptive field changes.

Gaussian filter width is defined by the full width at half maximum of the Gaussian profile, and is based on reported RGC dendritic field center sizes for the parasol, midget, and small bistratified RGCs [41, 42]. RGC receptive field overlap was determined according to literature as well, with neighbor distance set at their dendritic field diameter for midget cells, and half their dendritic field diameter for parasol and small bistratified cells [43].

At the output of the RGC convolution step, each unit represents the pooled response of a single retinal ganglion cell receptive field. To calculate relative contrast detection thresholds from the overall system, we used the probability summation of RGC signals to represent pooling by underlying cortical unit(s).

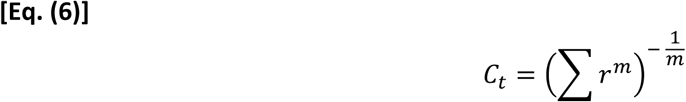

Here each RGC output (*r*) is raised to the probability summation exponent *m* then summed to achieve the final sensitivity of the cortical unit. The reciprocal of this cortical unit sensitivity provides relative detection thresholds. As the final goal of this simplified model is to simulate the spatial summation function, the relevant output is the inverse of system sensitivity, or relative detection thresholds as a function of stimulus size. To fully simulate the entire spatial summation function, stimuli of multiple sizes are passed through the model. For each stimulus size, 100 randomly selected positions were tested. While the photoreceptor and RGC lattice supposes a square array in our model, the results are not expected to differ significantly from that of hexagonal arrays [34].

## 3. RESULTS

### 3.1 Psychophysics

Spatial summation curves for all subjects and all conditions are shown in Figure 4, where log detection thresholds in cone contrasts are plotted as a function of log stimulus area in degrees squared. Here, different achromatic/chromatic conditions are color coded in rows, while eccentricities are separated in columns. For each subject, symbols represent the average thresholds for a specific stimulus size between all quadrants. Shown are the continuous hinged function fits for each individual, as well as the average of all subjects. Also shown are the model simulations under the same conditions. As simulation outputs are in relative threshold values, simulated curves were vertically scaled using the non-linear least squares method to fit the psychophysical data.

**Fig. 4.**
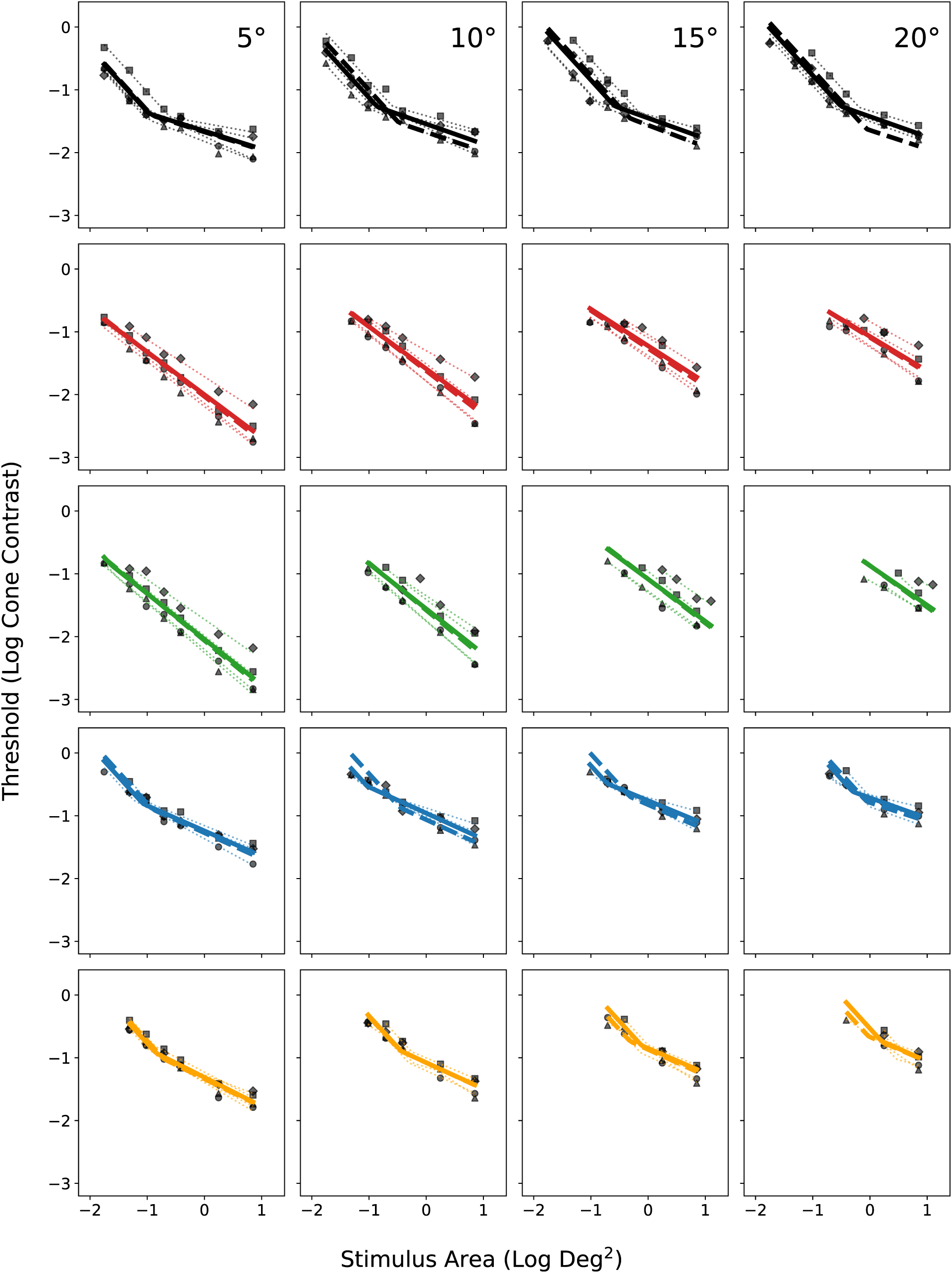
Continuous hinged functions obtained from different color conditions (rows) and eccentricities (columns) are plotted separately. Each subject’s results are represented by a specific symbol, showing the average threshold from the four quadrants for a given stimulus size. The smaller dotted line shows continuous hinged function fits for each individual. The larger solid line shows the mean of all individuals’ parameter values, while the larger dashed line shows the model simulations under the same conditions. In most cases, model fits overlap psychophysical averages and may be difficult to discern.

We chose to measure spatial summation functions at oblique meridians (quadrants) as sensitivity was not expected to vary significantly [29, 44] nor were we interested in meridional effects. To verify, a one-way ANOVA was conducted separately for each color, eccentricity, and stimulus size, resulting in 117 different conditions. This revealed no significant differences in thresholds between quadrants (p-value > 0.05), with the exception of two out of the 117 conditions. Latter results are reported for the average of the 4 meridians.

The continuous hinged function described the pooled psychophysics dataset well between all conditions (average R^2^ = 0.73). While the critical area was well characterized in the achromatic magnocellular, and chromatic koniocellular conditions, the chromatic parvocellular dataset lacked a distinct A_C_. For a closer look at the continuous hinged function fits to the psychophysical data, Figure 5 reveals changes in the *x*_0_, *y*_0_, and *b*_1_ parameters as a function of eccentricity for each of the color conditions. Following a similar convention as above, achromatic/chromatic conditions are color coded in rows, while individual parameters are separated in columns. Each subject’s average parameter values are shown for a given eccentricity as symbols, as well as the average of all subjects.

**Fig. 5.**
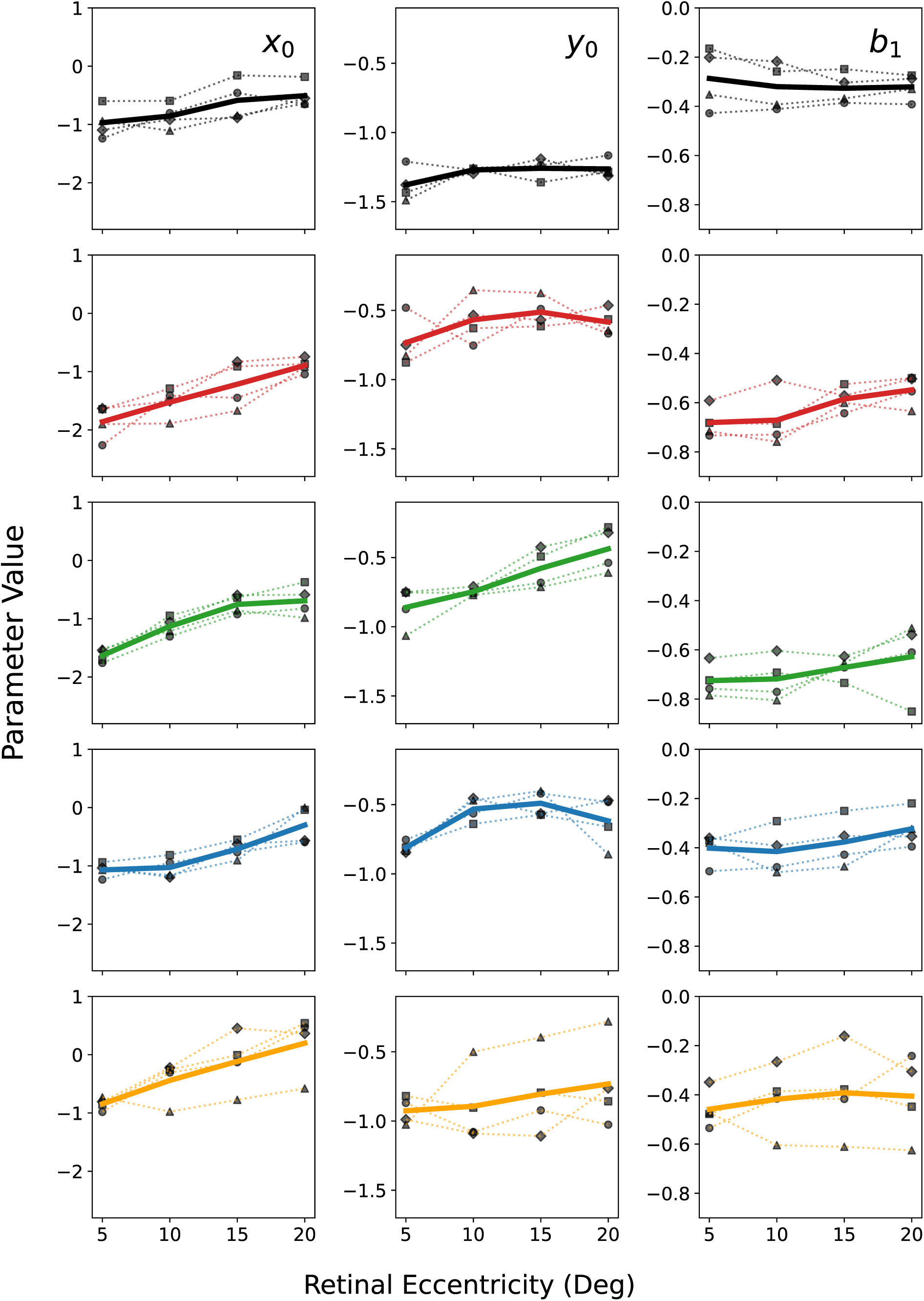
Average parameter values for the bootstrapped continuous hinged function fits shown for each chromatic condition (rows) and parameters (columns). *x*_0_, *y*_0_, and *b*_1_ parameters are plotted as a function of eccentricity. Each subject’s results are represented by a specific symbol along with a smaller dotted line, showing the average parameter value at a given eccentricity. The larger solid line shows the mean of all individuals. Due to the absence of complete spatial summation in the stimulus size range tested in chromatic parvocellular conditions (red & green), *x*_0_ parameter values follow the smallest stimulus size tested in each condition and represent the absolute upper bound. Likewise, the *y*_0_ parameter values follow the thresholds for the smallest stimulus size tested under parvocellular conditions and represent the lower bound.

As described in Equation 3, the *x*_0_ and *y*_0_ parameters refer to the x and y positions of the hinge of the continuous hinged function. This represents the critical area of complete spatial summation (*x*_0_) and the threshold at the A_C_ (*y*_0_). The *b*_1_ parameter refers to the slope of partial summation. As seen in Figure 4, chromatic parvocellular conditions (red & green) lacked areas of complete spatial summation, in conjunction with Figure 5, we see that the average *x*_0_ parameter values follow the smallest stimulus size tested. Due to the lack of data for small enough stimulus sizes to characterize areas of complete spatial summation under these conditions, the *x*_0_ and *y*_0_ parameter values are not well pinned down. By extension, the respective *y*_0_ parameter follows the threshold for the smallest stimulus size tested.

A one-way ANOVA conducted separately for each eccentricity revealed a significant difference in all parameter values between chromatic conditions (all p-values < 0.05). Further, a one-way ANOVA conducted separately for each chromatic condition revealed an effect of eccentricity on all parameter values (all p-values < 0.05). On a more qualitative note, achromatic magnocellular conditions are the most similar to chromatic koniocellular conditions at least for the *x*_0_ and *b*_1_ parameter values. This is seen in Figure 5 where critical areas and the slope of partial summation align quite well at a given eccentricity.

### 3.2 Receptive Field Pooling Model

To understand how the A_C_ depends on responsible visual processing pathway, the RGC convolutional filtering level of the model is split into separate retinal ganglion cell types, the parasol cells, midget cells, and small bistratified cells. These cell types with distinct dendritic field diameters determine the size of the filtering kernel or receptive field at this level. To understand the effects of retinal eccentricity on the A_C_ within these different pathways, the filtering kernel size was adjusted according to changes in dendritic field size with eccentricity [41, 42]. As small bistratified dendritic field size matches parasol cell size closely within the central 20°, the model reflects just the parasol and midget cells (i.e. parasol dendritic field sizes were used for small bistratified calculations).

Figure 6 summarizes the effects of retinal receptive field size and the probability summation exponent *m* on simulated spatial summation functions. Retinal receptive field sizes shown are derived from parasol retinal ganglion cell dendritic field sizes as a function of retinal eccentricity. Here, the continuous hinged function was fit to the model outputs using the non-linear least squares method, providing the *x*_0_, *y*_0_, and *b*_1_ terms. RGC dendritic field diameter is shown to be directly proportional to the A_C_, while having no effect on the slopes of the complete and partial summation areas. As a consequence of summation, decreasing receptive field size decreases thresholds. As each RGC unit contributes equally to detection, decreasing the size of the receptive field increases their density and thus the number of contributing receptive fields, improving thresholds. On the right panel, the probability summation exponent *m* is shown to primarily affect the slope of partial summation.

**Fig. 6.**
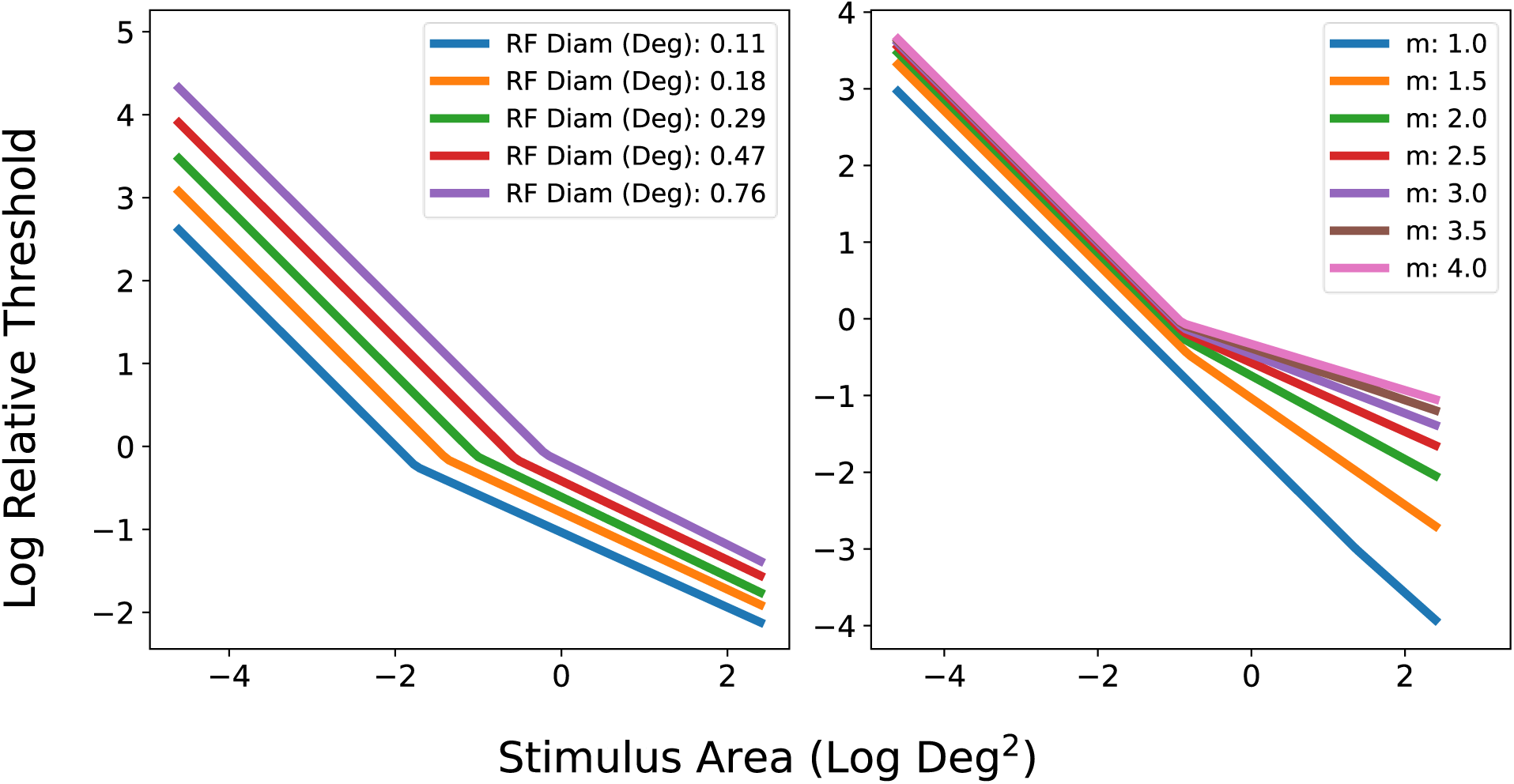
**(left).** Shown are example model simulations given 5 different retinal eccentricities. Receptive field (RF) diameters are calculated for parasol ganglion cell dendritic fields as a function of retinal eccentricity. Here, receptive field size is shown to be directly related to the critical area. **(right).** Shown are example model simulations given 6 different probability summation *m* exponent values. Here, the slope of partial summation (*b*_1_) is shown to be inversely related to the *m* exponent.

Model simulations are overlaid on the psychophysical data in Figure 4 (dashed lines). Spatial summation functions were simulated for each of the chromatic conditions and for each eccentricity tested. As literature on the slopes of partial summation is scarce, particularly within our targeted pathways, we derived them from empirical fits to the psychophysical data, and defined a *b*_1_ parameter to *m* exponent conversion factor. Interestingly the *m* exponent changes both as a function of eccentricity [45] and cell type. The model outputs were fit with the continuous hinged function to determine the critical area (*x*_0_), and slope of partial summation (*b*_1_). As the model threshold outputs are arbitrary, the continuous hinged function with fixed *x*_0_ and *b*_1_ parameters were again fit using the non-linear least squares method to the psychophysical data to determine a proper *y*_0_ offset value.

Our receptive field pooling model based on retinal ganglion cell dendritic field size calculated from respective RGC type and retinal eccentricity were able to predict the critical area and match psychophysical functions (average R^2^ = 0.71). In the achromatic condition, the average *m* parameter value ranged from 3.8 – 4.5, decreasing with eccentricity. On the other hand, the *m* parameter in chromatic conditions all increased with eccentricity. In the parvocellular red condition, the average *m* parameter value ranged from 1.4 – 1.8. In the parvocellular green condition, the average *m* parameter value ranged from 1.3 – 1.5. In the koniocellular blue condition, the average *m* parameter value ranged from 2.7 – 3.9. In the koniocellular yellow condition, the average *m* parameter value ranged from 2.4 – 3.0. This *m* exponent of probability summation was able to account for the variety of partial summation slopes in the psychophysical results.

## 4. DISCUSSION

In the present study, we measured changes in visual sensitivity within targeted magnocellular, parvocellular, and koniocellular pathways as a function of stimulus size and retinal eccentricity. We also describe a physiologically motivated model that can account for differences in summation characteristics between these pathways and within the eccentricities tested. Figure 4 reveals the simulated spatial summation functions accompanied by psychophysically measured curves. Our simple model based on the probability summation of early receptive field units was able to account for the spatial summation functions of different visual processing pathways and across eccentricities.

Figure 7 provides a summary of the continuous hinged function fits to the pooled psychophysical data. All chromatic conditions are shown together for each of the eccentricities tested. Each curve is truncated at the minimum and maximum stimulus sizes tested. For all chromatic conditions, the lower stimulus size bound is the smallest size detectable with our current setup. Also shown are the Goldmann sizes for standard automated perimetry. While complete and partial summation areas are well characterized for the magnocellular (achromatic) and koniocellular (blue/ yellow) conditions, parvocellular (red/ green) conditions lacked a critical area for the stimulus size range tested. Instead, the data seem to be best fit by a single linear function. As the slope of these lines are shallower than the −1 slope expected for complete summation, it is likely that the data set reveals partial summation areas. Non-linear least squares fitting with the continuous hinged function suggests this as well, where the bootstrap distribution for the *x*_0_ parameter lies to the left of the smallest stimulus size tested in the parvocellular conditions.

**Fig. 7.**
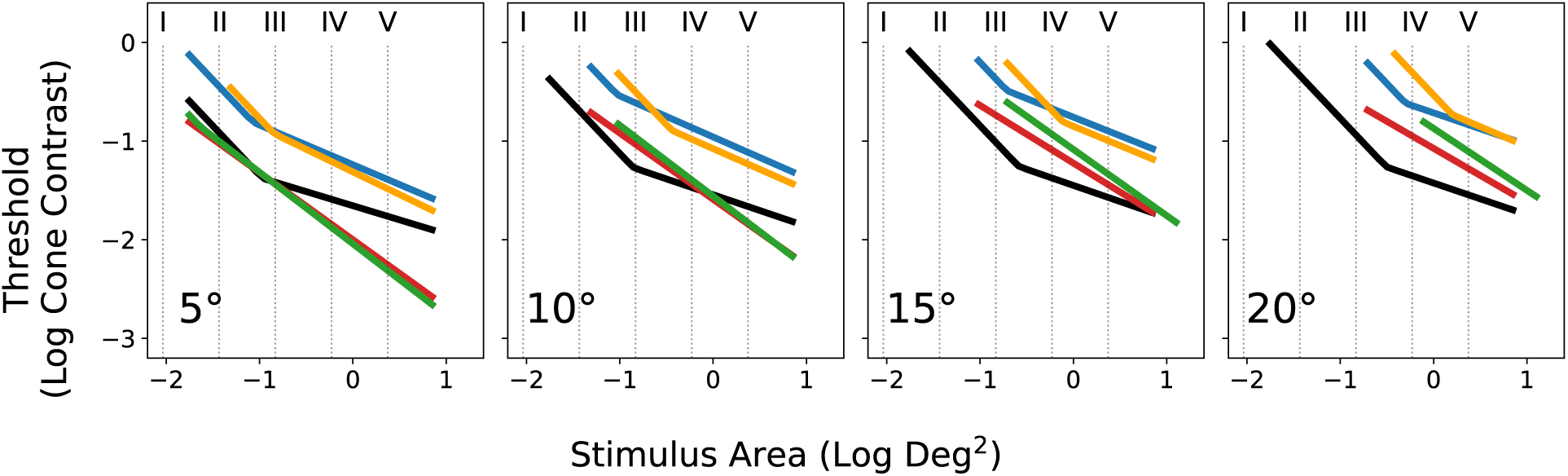
Plot of psychophysical results for each color condition, and each eccentricity tested (columns). Goldmann sizes for standard automated perimetry are overlayed for reference.

There has been great interest in tying the psychophysical measure of the critical area to some neurological mechanism or anatomical structure. The idea that the A_C_ is associated with the RF centers of RGCs has been considered previously in studies limited to achromatic contrast. Recent studies involving more specific chromatic stimuli have considered the density or number of underlying RGCs [9, 10]. While this has been thought to hold true for achromatic and koniocellular stimuli, inconsistencies exist and fail to explain parvocellular findings [25]. Despite these attempts to link the A_C_ to some aspect of the visual system, most fail to account for previously reported changes in the A_C_ as a result of non-spatial stimulus features. For example, the A_C_ has been shown to shrink with increasing background luminance [46], raising the difficulty of associating it with some set anatomical structure rather than some physiological characteristic. In this way, it has been suggested that A_C_ is regulated to maintain contrast thresholds for stimuli within a certain size regardless of background intensity [5]. It has also been suggested that the A_C_ shifts to maintain a constant Weber fraction [6].

Figure 8 provides a summary of the average bootstrapped continuous hinged function fit parameters as a function of retinal eccentricity. Here, the *x*_0_ parameter represents the critical area, while the *b*_1_ parameter represents the slope of partial summation. For the *x*_0_ parameter, parvocellular (red/ green) data was omitted as complete spatial summation was lacking for the stimulus size range tested. Instead, the smallest stimulus size tested for these conditions is marked (stars). Also shown are average parasol (magnocellular), midget (parvocellular), and small bistratified (koniocellular) retinal ganglion cell dendritic field diameter estimates from literature [41, 42]. Small bistratified RGC dendritic field sizes have been found to be quite similar to that of the parasol RGCs [42] and are shown together.

**Fig. 8.**
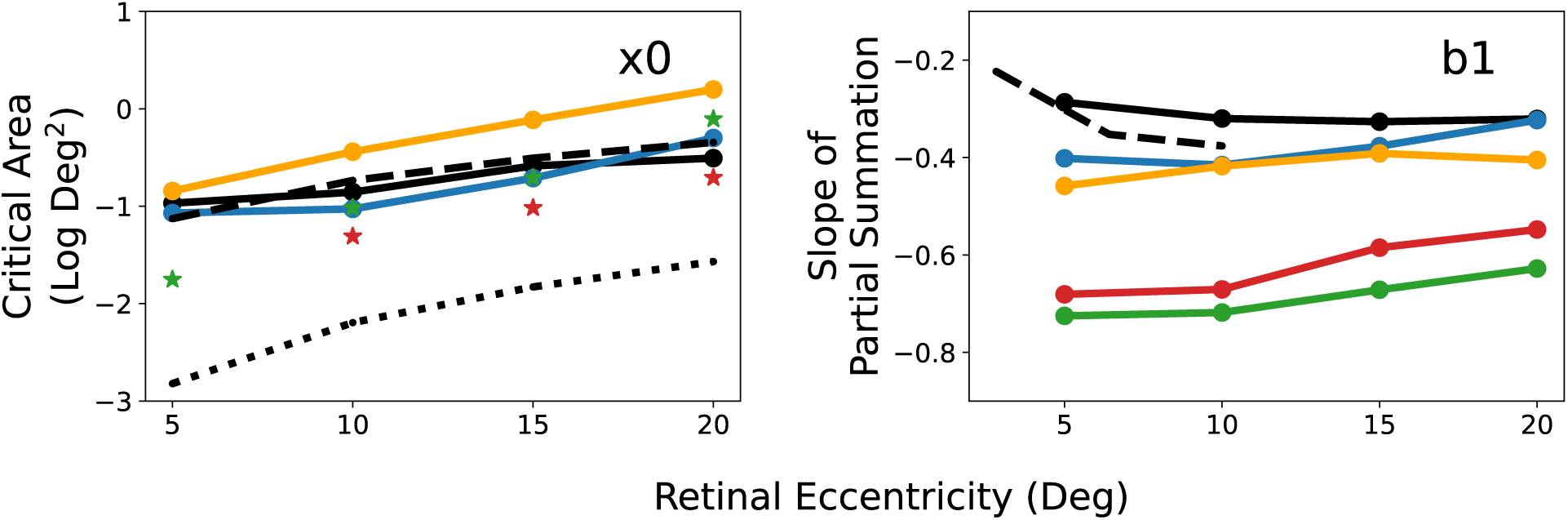
**(left).** Plot of the continuous hinged function *x*_0_ parameter, representing the critical area. As psychophysical data lacked small enough stimulus sizes to adequately characterize the A_C_ for parvocellular (red/ green) conditions, they were omitted. Instead, the smallest stimulus sizes tested are shown (red/ green stars). Also shown are average dendritic field diameter measures from literature for magnocellular and koniocellular conditions (dashed line) and parvocellular conditions (dotted line). **(right).** Plot of the continuous hinged function *b*_1_ parameter, representing the slope of partial summation. The dashed line represents achromatic partial summation slopes averaged from oblique meridians of 10-2 standard automated perimetry test format from literature [45].

The parasol and small bistratified RGC dendritic field size aligns well with achromatic and koniocellular critical area estimates. In these conditions, the A_C_ increased monotonically within the eccentricities tested. The midget RGC dendritic field size however, is much smaller than that of the parasol and small bistratified RGCs, and much smaller than the smallest size we were able to test in the parvocellular conditions. The smallest stimulus sizes tested are more than a log unit larger than their respective midget RGC dendritic field sizes. The lack of critical area in the chromatic parvocellular condition is theorized to be a consequence of the smaller midget RGC sizes and our stimulus size ranges. While it is natural to test smaller stimulus sizes in the parvocellular condition to capture complete summation areas and the critical area, our subjects were unable to perceive stimuli smaller than the sizes tested in our current experimental setup. It is possible that decreasing overall luminance could enhance the detectability of the targets and increase the A_C_ to a perceivable range [46]. However, it is unknown whether the increase in A_C_ with decreasing luminance would hold under chromatic conditions.

Previous studies on the spatial summation characteristics of chromatic contrasts specified by the cardinal axes of DKL color space report similar difficulties in determining contrast detection thresholds for smaller chromatic stimuli, especially for the “green” mechanism [25]. Asymmetries between the increment and decrement directions of the parvocellular condition (red/ green), are difficult to extrapolate from our results as we were unable to characterize the A_C_ for these conditions. However, asymmetries between the increment and decrement directions of the koniocellular condition (blue/ yellow) follow expectations from literature with “blue” critical areas being slightly smaller than that of “yellow” [10, 25]. Rather than asymmetries between opponent directions of the same chromatic axes, our results show greater difference between parvocellular functions compared to that of the koniocellular or magnocellular pathways. Nevertheless, asymmetries in performance for opponent colors within a mechanism are well documented [47], as are performance differences between different visual processing mechanisms [48].

Aside from chromatic methods to isolate post-receptoral visual mechanisms, there is extensive literature on the use of achromatic stimuli with specific temporal characteristics to target the same mechanisms. The steady and pulsed pedestal paradigms [48, 49] take advantage of the different temporal processing characteristics of the magnocellular and parvocellular pathways as a method of isolation. Despite differences in isolation methods, the shape of spatial summation functions within the targeted magnocellular and parvocellular pathways using the steady and pulsed pedestal paradigms [50, 51] are quite similar to those obtained in our current study. In those studies, magnocellular functions were fit with a continuous decay function, which was able to capture the steepness of complete summation areas and shallowness of partial summation areas. Similar to our findings, parvocellular data were better fit with a linear function within the size range tested and presented with a slope shallower than −1.

Figure 8 (right) provides a summary of the continuous hinged function *b*_1_ parameter (representing the slope of partial summation) as a function of retinal eccentricity. The dashed line represents achromatic partial summation slopes averaged from the oblique meridians of the 10-2 standard automated perimetry test format [45]. Eccentricities are calculated from the square grid pattern of the 10-2 test array. While we lack data within the central 5 degrees, we do capture a similar steepening of the achromatic partial summation slope with eccentricity. As with the critical area, different chromatic conditions/ mechanisms reveal distinctly different slopes and profiles. Parvocellular and koniocellular conditions become shallower with eccentricity, with parvocellular slopes being much steeper. It is unknown what the slope of partial summation reflects in terms of visual mechanism characteristics.

Figure 9 shows an alternative visualization of the data, with thresholds in cone contrast plotted as a function of retinal eccentricity for four stimulus sizes in columns. As with previously discussed parameters, there is a clear difference in performance between the magnocellular, parvocellular, and koniocellular conditions. Magnocellular thresholds were relatively insensitive to eccentricity, as described in literature [23, 45]. Chromatic thresholds on the other hand, show a greater rate of change with eccentricity. Near the fovea, parvocellular thresholds were smaller than that of the magnocellular and koniocellular pathways. This is in agreement with previous reports of greater foveal sensitivity for chromatic targets compared to achromatic targets [52]. While thresholds were quite similar between opponent directions along the same chromatic mechanism, certain asymmetries exist here as well. In both the decremental parvocellular (green) and decremental koniocellular (yellow) conditions, smaller targets were difficult to detect in the periphery. There have been extensive discussions on the difficulty in detecting smaller green targets [25, 47].

**Fig. 9.**
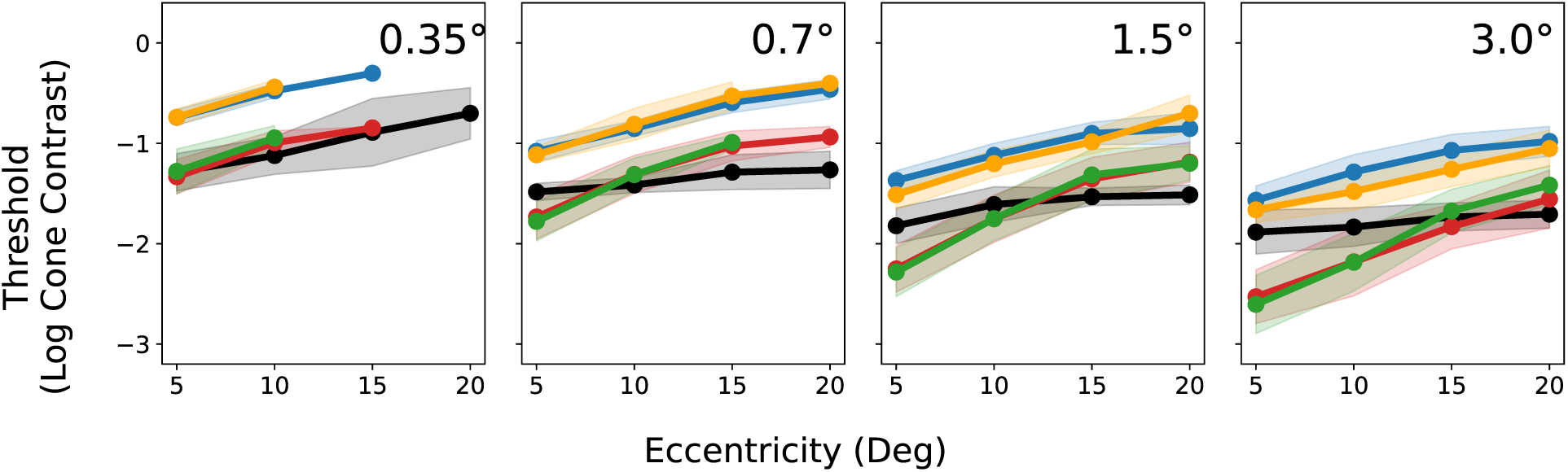
Chromatic detection thresholds in cone contrast as a function of retinal eccentricity for 4 different stimulus sizes (columns, 0.35°, 0.7°, 1.5°, 3.0° diameter). Markers are the average of all subjects and positions tested for a given color condition and eccentricity, with error bars revealing one standard deviation from the mean. Certain conditions may have fewer subjects capable of perceiving the target, shown by an absent point or the lack of an error bar.

This alternative perspective of the data reveals a limitation of our model, in that it is unable to replicate these particular nuances or asymmetries in human performance. Nevertheless, our model, is able to account for differences in the critical area between pathways, as well as its changes with eccentricity. We have also shown that the −1 slope of complete summation and the shallower slope of partial summation areas can be accounted for through the probability summation of retinal receptive field units. While alternative models that describe spatial summation exist [19, 20, 21, 53], our goal was to develop a simple model that could account for differences in the spatial summation function between visual processing pathways and across eccentricities. Pan & Swanson [19] described the probability summation of multiple spatially tuned cortical units, and were able to account for a variety of findings in literature across eccentricities. Meese & Summers [20, 21] on the other hand, introduced an extra stage of internal additive noise to their model to account for their data. Montesano *et al.* [53] proposed a spatiotemporal integration model based on the convergence of cone photoreceptors to underlying retinal ganglion cells to account for both the spatial and temporal summation function. Since we were primarily interested in the differences between processing pathways, we chose to focus on retinal ganglion cell characteristics with idealized cortical pooling using probability summation. It is possible that the merging of these models could yield even greater insights about the impacts of physiological properties on detection thresholds.

Our model suggests that the critical area of summation is driven by retinal ganglion cell receptive field size, while the slope of partial summation can be controlled by the *m* exponent of probability summation. We hypothesize that while a stimulus is within the receptive field of an RGC, complete summation occurs. As the stimulus exceeds the size of that receptive field, neighboring fields begin to contribute to its detection, here modeled by probability summation. This detection is modulated by numerous factors, but can be approximated by the probability summation exponent *m*. Understanding these factors can provide an explanation for differences in partial summation slopes found between studies in literature, between eccentricities [45], and visual processing pathways.

In our study, the *m* exponent range between 3.8 and 4.5 under achromatic magnocellular conditions is slightly higher than previous estimates between 3 and 4 [35, 54], however the contemporary use of 4 is well within our calculated range [19, 53, 55]. Interestingly *m* estimates under the koniocellular blue condition (2.7-3.9) match the 3-4 range found in literature, while the koniocellular yellow condition revealed slightly smaller *m* estimates (2.4-3.0). Similar asymmetries exist with the *m* estimates of the parvocellular condition, with the red condition (1.4-1.8) being slightly larger than that of the green condition (1.3-1.5). Parvocellular *m* estimates were much smaller than that of the magnocellular or koniocellular. It is unknown what the *m* exponent reflects in terms of physiological differences in processing between pathways, although it is likely governed by higher level cortical mechanisms.

In summary, we have revealed that the critical area of complete spatial summation aligns with the dendritic field size of underlying retinal ganglion cells within the central 20 degrees of visual field, for the magnocellular and the koniocellular pathways. For the parvocellular pathway, however, we found only partial summation within a similar stimulus size range. We hypothesize that this is due to the exceptionally small size of midget retinal ganglion cell dendritic fields. An anatomically and physiologically motivated model of contrast pooling through probability summation was able to account for these differences in spatial summation between mechanisms, and also describe changes in the critical area with retinal eccentricity.

